# Engineering single pan-specific ubiquibodies for targeted degradation of all forms of endogenous ERK protein kinase

**DOI:** 10.1101/2021.04.16.440242

**Authors:** Erin A. Stephens, Morgan B. Ludwicki, Bunyarit Meksiriporn, Mingji Li, Tianzheng Ye, Connor Monticello, Katherine J. Forsythe, Lutz Kummer, Andreas Plückthun, Matthew P. DeLisa

## Abstract

Ubiquibodies (uAbs) are a customizable proteome editing technology that utilizes E3 ubiquitin ligases genetically fused to synthetic binding proteins to steer otherwise stable proteins of interest (POIs) to the proteasome for degradation. The ability of engineered uAbs to accelerate the turnover of exogenous or endogenous POIs in a posttranslational manner offers a simple yet robust tool for dissecting diverse functional properties of cellular proteins as well as for expanding the druggable proteome to include tumorigenic protein families that have yet-to-be successfully drugged by conventional inhibitors. Here, we describe the engineering of uAbs comprised of a highly modular human E3 ubiquitin ligase, human carboxyl terminus of Hsc70-interacting protein (CHIP), tethered to different designed ankyrin repeat proteins (DARPins) that bind to nonphosphorylated (inactive) and/or doubly phosphorylated (active) forms of extracellular signal-regulated kinase 1 and 2 (ERK1/2). Two of the resulting uAbs were found to be global ERK degraders, pan-specifically capturing all endogenous ERK1/2 protein forms and redirecting them to the proteasome for degradation in different cell lines, including MCF7 breast cancer cells. Taken together, these results demonstrate how the substrate specificity of an E3 ubiquitin ligase can be reprogrammed to generate designer uAbs against difficult-to-drug targets, enabling a modular platform for remodeling the mammalian proteome.

## Introduction

Proteome editing technology represents a powerful strategy for posttranslational control of protein function based on the principle of “inhibition-by-degradation” whereby an inhibitor/degrader hijacks the cellular quality control machinery to selectively eliminate target proteins ^1–3^. A common feature of proteome editing approaches is the ability to promote catalytic turnover of otherwise stable intracellular proteins, requiring only transient binding to virtually any site on the protein of interest (POI). This is in stark contrast to traditional occupancy-based inhibitors, which depend on a distinct binding site that affects function (*e.g*., enzyme active site) and require relatively high concentrations to ensure sustained stoichiometric binding. For these reasons, the development of proteome editors that are capable of inducing protein degradation is gaining considerable attention for both scientific investigation of native protein function and therapeutic targeting of disease-relevant proteins, especially those that are recalcitrant to conventional pharmacological interventions and have thus been deemed difficult-to-drug ^4, 5^.

The creation of customized degrader molecules typically involves precision marking of specific POIs for proteolytic removal via molecular mimicry of natural degradation processes found in eukaryotic cells. The most frequently exploited of these degradation processes is the ubiquitin-proteasome pathway (UPP), which involves the sequential activities of three enzymes – ubiquitin-activating enzyme (E1), ubiquitin-conjugating enzyme (E2), and ubiquitin ligase (E3) – that cooperate in an energy-dependent manner to covalently tag available protein lysines with a polyubiquitin chain ^6^. While a variety of polyubiquitin chain topologies are possible, K48-linked ubiquitin serves as the canonical recognition signal for the 26S proteasome and generally leads to substrate degradation ^7^. The fact that E3s govern substrate specificity and often exhibit remarkable plasticity has made these enzymes the component of choice in the majority of proteome editing technologies described to date. Most notable among these technologies is PROTACs (proteolysis targeting chimeras) ^8, 9^, which are heterobifunctional small molecules that effectively bridge the E3 and the POI, forming a ternary complex that triggers target polyubiquitination and subsequent proteasomal degradation in cultured cells and mice ^10–14^. With respect to clinical potential, two PROTACs, named ARV-110 and ARV-471, targeting androgen receptor and estrogen receptor, respectively, have advanced into phase I human trials ^15^.

Alongside small-molecule PROTACs are protein-based chimeras in which an E3 is genetically fused to a peptide or protein with affinity for the POI. In the earliest designs, substrate targeting was achieved by leveraging naturally occurring protein interaction partners, whereby fusion of an E3 (or a component of an E3 ligase complex) to a POI’s known binding partner yielded a chimera that promoted knockout of the cognate POI following expression in cultured cells ^16, 17^. When a binding partner for a given POI is available, this approach has proven to be highly effective both *in vitro* and *in vivo*, leading to induced degradation of several different oncoprotein targets including c-Myc, ErbB, HIF-α, and KRAS ^18–21^. However, this approach is limited to only those POIs for which a natural interacting partner is known.

Therefore, to extend this approach beyond naturally occurring protein-protein interactions, we created ubiquibodies (uAbs) by genetically fusing an E3 to a synthetic binding protein such as a single-chain antibody fragment (scFv), a designed ankyrin repeat protein (DARPin), or a fibronectin type III (FN3) monobody ^22^. Because synthetic binders can be readily identified using methods such as phage, ribosome, and yeast display ^23, 24^ with the potential for proteome-scale coverage ^25^, uAbs are a universally applicable technology that can be developed against virtually any intracellular POI. Indeed, by combining the flexible ubiquitin-tagging capacity of a human RING/U-box-type E3 named CHIP (carboxyl-terminus of Hsc70-interacting protein) with the programmable affinity and specificity of synthetic binding proteins, we demonstrated that uAbs efficiently redirected *Escherichia coli* β-galactosidase (β-gal) and maltose-binding protein (MBP) to the UPP for proteolytic degradation ^22^. Importantly, neither of the POIs was a natural substrate for CHIP and the degradation that we observed did not depend on the biological function or interaction partners of the POIs. Also noteworthy is the highly modular architecture of uAbs: swapping synthetic binding proteins enables generation of new uAbs that recognize completely different POIs ^26–34^ while swapping E3 domains enables tailoring of the catalytic efficiency and/or E2 specificity ^27, 34^. It is even possible to deplete certain protein subpopulations (*e.g*., active/inactive, posttranslationally modified/unmodified, wild-type (wt)/mutant, *etc*.) while sparing others ^17, 18, 27^.

Here, we exploited the versatility of uAbs to construct proteome editors capable of selectively removing the major isoforms of extracellular signal-regulated kinase (ERK), namely ERK1 and ERK2, which share ~85% identity in their amino-acid sequence and appear to be functionally equivalent ^35^. Following activation by phosphorylation on tyrosine and threonine residues by upstream kinases in the mitogen-activated protein kinase (MAPK) pathway ^36^, ERK1/2 phosphorylate numerous substrates that participate in key physiological processes that control cell proliferation, differentiation, survival, and death ^37, 38^. We chose to focus on ERK1/2 because the MAPK pathway is the most frequently mutated signaling pathway in human cancer, making components of this cascade attractive targets for drug development ^36^. To this end, a significant number of RAF and MEK inhibitors have been preclinically and clinically evaluated, which is in contrast to the more limited development of selective ERK1/2 inhibitors. While there are many reasons for this discrepancy ^36^, occupancy-based inhibitors specific for ERK are very difficult to design due to the high homology between active-site pockets of ERK1/2 and cyclin-dependent kinases (CDKs).

To address this challenge, we generated a global ERK degrader by recombining human CHIP’s discrete catalytic U-box domain with a pan-specific DARPin named EpE89 that recognizes both nonphosphorylated ERK1 and ERK2 as well as the doubly phosphorylated forms, pERK1 and pERK2 ^39^. Our results demonstrated the efficacy of this engineered uAb, as well as a second design based on a pERK1/2-specific DARPin named pE59 ^39^, in pan-selectively inducing ubiquitin-mediated degradation of all major ERK1/2 proteoforms in cultured cells. In addition, we uncovered the molecular basis for pan-specificity, which appeared to originate from an ability of the engineered uAbs to install polyubiquitin, including K-48-linked chains, on both ERK2 and pERK2.

## Results

### Construction of a pan-specific ERK ubiquibody

CHIP is a human U-box E3 ubiquitin ligase with three discrete domains, an N-terminal tetratricopeptide repeat (TPR) domain, a C-terminal U-box domain, and an intermediate coiled-coil linker (**Fig. 1a**) ^40, 41^. The TPR domain of CHIP binds to the molecular chaperones Hsc70-Hsp70 and Hsp90, facilitating ubiquitination of chaperone-bound client proteins ^42^. To convert CHIP into a pan-specific ERK degrader, we replaced its N-terminal TPR domain with DARPin EpE89 that recognizes nonphosphorylated and doubly phosphorylated ERK1 and ERK2 (**Fig. 1a**) ^39^. For comparison purposes, we constructed two additional uAbs comprised of phospho-isoform-specific DARPin pE59, which preferentially binds pERK1 and pERK2, and DARPin, E40, which specifically recognizes nonphosphorylated ERK1 and ERK2. Our uAb designs retained the flexible coiled-coiled domain of CHIP, which has been shown to be critical for E3 dimerization ^40^, as well as the catalytic U-box domain. A short linker of five amino acids (Gly-Ser-Gly-Ser-Gly) was included to ensure flexibility between the C-terminal capping helices of the DARPin and helix α7 of the N-terminally truncated CHIP (CHIPΔTPR) (**Fig. 1a and b**). The rationally designed uAbs were expressed in the cytoplasm of *E. coli* cells and purified by Ni-NTA affinity chromatography, resulting in soluble titers (~30 mg protein per liter culture) that were notably higher than their unfused DARPin counterparts (**Supplementary Fig. 1a**). This latter observation indicated that the CHIPΔTPR domain somehow enhanced the expression of its DARPin fusion partners. Following purification and characterization by size exclusion chromatography (SEC), the uAbs were observed to elute slightly earlier than the non-aggregated portion of wild-type CHIP (**Supplementary Fig. 1b**). Since CHIP eluted at a volume expected of a dimer with a large water shell, consistent with the observation that the U-box of human CHIP functions as a homodimer ^40, 41^, we concluded that the uAbs were similarly assembled as dimeric structures akin to their parental E3 ubiquitin ligase.

**Figure 1.**
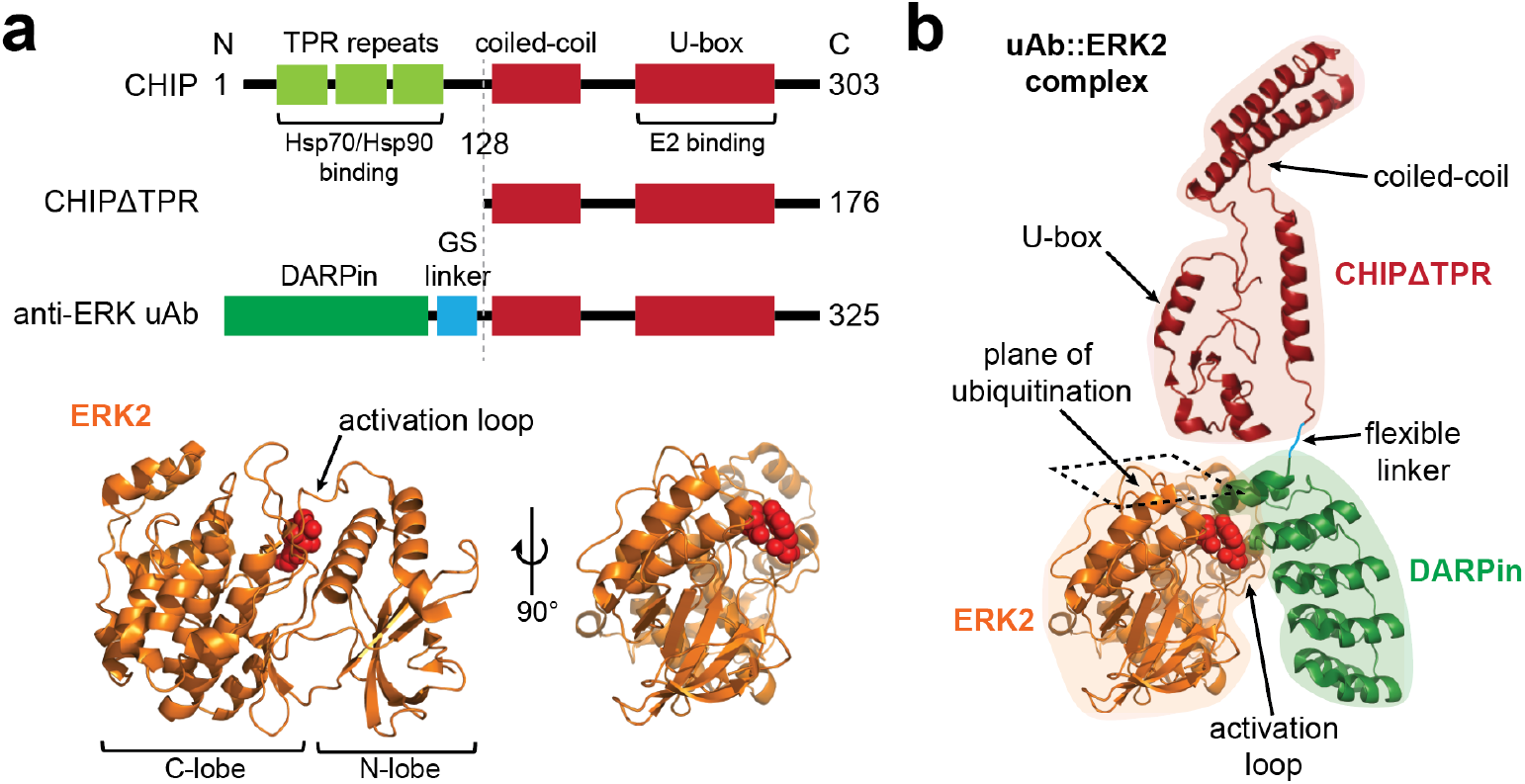
Modular design of a pan-specific ERK degrader. (a) The architecture of uAbs is highly modular, involving three distinct domains: a substrate-binding domain comprised of a synthetic binding protein (green); a flexible Gly-Ser-Gly-Ser-Gly (GS) linker (blue); and a catalytic domain comprised of the C-terminus of human CHIP starting from residue 128 (CHIPΔTPR) (red). The synthetic binding domains used in this study are DARPins with specificity for different ERK forms. Crystal structure of human ERK2 (orange) generated in PyMOL (PDB ID: 3ZU7). Residues T185 and Y187 (red balls) are phosphorylated upon activation, leading to rotation of the N-lobe relative to the C-lobe. (b) Orientation of the catalytic domain in uAb::ERK2 complex. The ubiquitination plane is in direct proximity to the theoretical position of the catalytic domain. Schematic generated from a composite of PDB ID: 2C2L and 3ZU7 in PyMOL and Illustrator software.

### Reprogramming the substrate specificity of CHIP with ERK-binding DARPins

The extent to which CHIP’s substrate specificity was switched by tethering to pan-ERK-specific EpE89 was first evaluated using a previously described affinity precipitation assay ^39^. In this assay, lysate derived from human embryonic kidney (HEK) 293T cells, a common epithelial cell line, was incubated with purified EpE89-uAb, which was subsequently captured by Ni-NTA beads. Immunoblotting analysis revealed that EpE89-uAb was able to precipitate endogenous ERK1 and ERK2 as evidenced by the cross-reactivity of elution fractions with a phosphorylation-state independent anti-ERK antibody that recognizes all ERK isoforms including phosphorylated ones (**Supplementary Fig. 2a**). Similar affinity precipitation was achieved with pE59-uAb, E40-uAb, and the unfused DARPins, with the behavior of the latter in agreement with Kummer *et al* ^39^. In contrast, CHIPΔTPR and the non-specific control Off7-uAb, a chimera between CHIPΔTPR and the DARPin Off7 that binds *E. coli* maltose-binding protein ^43^, were unable to capture ERK1/2. Importantly, none of the proteins precipitated Hsp70, a native substrate of full-length CHIP ^42^, indicating that CHIP’s substrate specificity had been effectively reprogrammed by swapping the TPR domain with ERK-binding DARPins.

To evaluate the pan-specificity of EpE89-uAb in more detail, we performed an enzyme-linked immunosorbent assay (ELISA) using ERK2 and pERK2 as immobilized antigens. Consistent with the known binding specificity of unfused EpE89 ^39, 44^, the EpE89-uAb bound avidly to both ERK2 and pERK2. The pE59-uAb and E40-uAb constructs similarly mirrored the substrate preferences of their parental DARPins, specifically binding pERK2 and ERK2, respectively, at levels that rivaled the binding activity of pan-specific EpE89-uAb for each target (**Fig. 2a** and **Supplementary Fig. 2b**). It should be noted that while pE59-uAb and its unfused pE59 counterpart clearly preferred cognate pERK2, each bound to nonphosphorylated ERK2 at a low but reproducible level above background. A similar pattern was observed for E40-uAb and E40, with each preferring ERK2 but showing a low level of binding to non-cognate pERK2. These results are consistent with previous findings that the binding affinity between each of these DARPins and its non-cognate ERK2 or pERK2 form, while significantly weaker than with the cognate form, were still in the low micromolar range ^39^. Importantly, the N-terminally truncated CHIPΔTPR construct, which lacked a substrate-binding domain, showed no measurable binding activity above background to either ERK2 or pERK2 (**Fig. 2a** and **Supplementary Fig. 2b**). The enhanced binding measured for the dimeric uAbs relative to the unfused DARPins is likely due to an avidity effect, as the uAbs are dimers wheras DARPins are monomeric. Overall, these results indicate that the DARPins successfully reprogrammed CHIP specificity for distinct ERK forms, with EpE89-uAb showing the clearest capacity for pan-specific ERK silencing.

**Figure 2.**
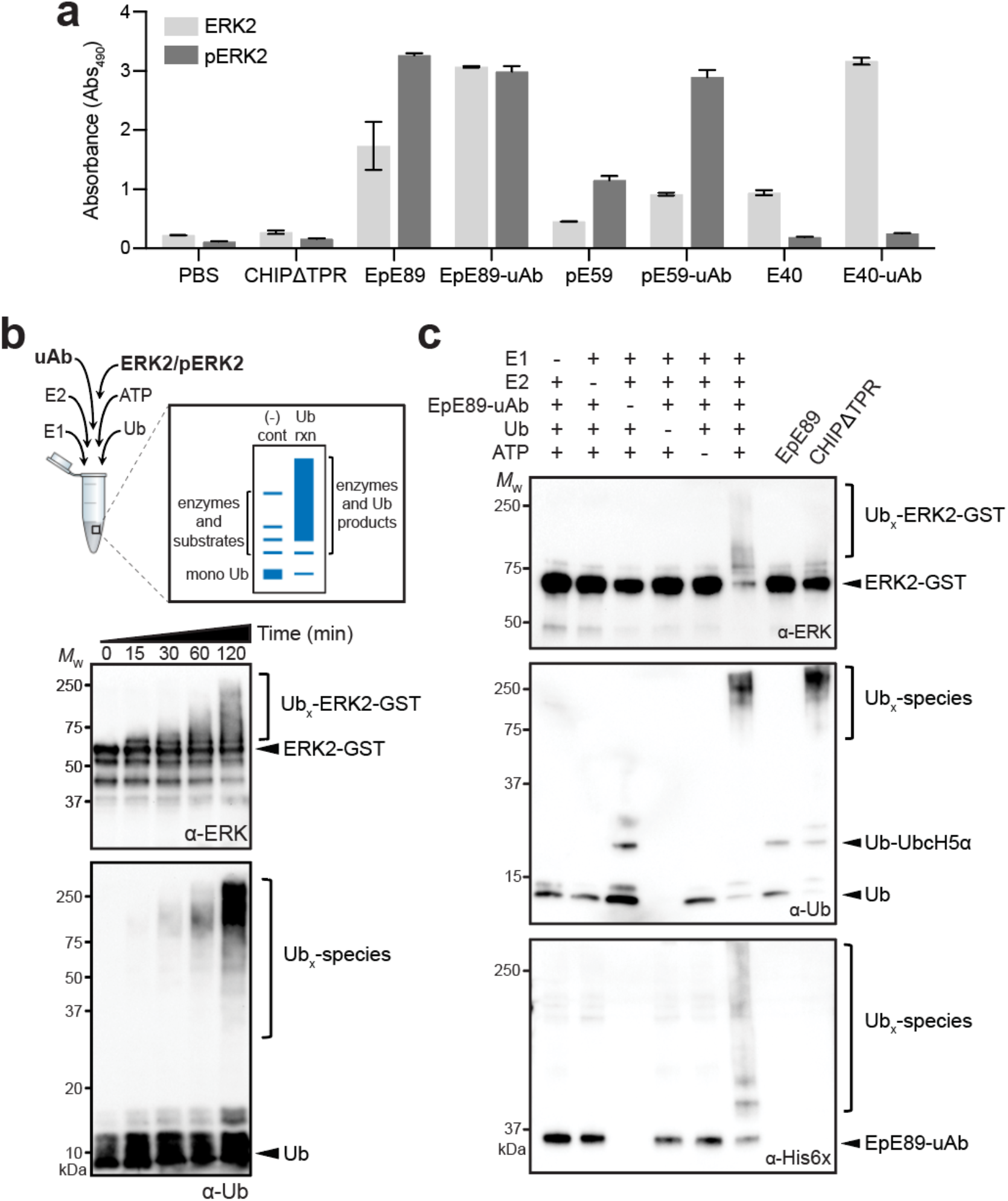
Engineered uAbs bind and ubiquitinate ERK *in vitro*. (a) ELISA of purified uAbs, DARPins, and CHIPΔTPR against immobilized ERK2 or pERK2 as indicated. Buffer only (PBS) served as a negative control. An equivalent amount of each uAb, DARPin, and CHIPΔTPR protein was used in the assay. Data are average of three biological replicates and error bars represent standard deviation of the mean. (b) *In vitro* ubiquitination of nonphosphorylated ERK2 (ERK2-GST) in the presence of purified EpE89-uAb along with E1, E2, ubiquitin (Ub), and ATP. Samples were collected at indicated times and subjected to immunoblotting. (c) Same as in (b) but all reactions were run for 120 min in presence (+) or absence (-) of each pathway component as indicated. Controls included EpE89 and CHIPΔTPR in presence of all pathway components. For all blots, an equivalent amount of total protein was added to each lane. Immunoblots were probed with: pan-ERK antibody (α-ERK) and anti-ubiquitin (α-Ub) to detect ERK2 and Ub species, respectively; and anti-His6x antibody to detect uAb. Protein bands corresponding to ERK2, Ub, EpE89-uAb, and polyubiquitinated species (Ub_x_) are marked at right. Molecular weight (*M*_W_) markers are indicated at left. Results are representative of at least three biological replicates.

### Ubiquibodies promote ubiquitin transfer to ERK

Having demonstrated that EpE89-uAb possessed pan-specific ERK binding, we next performed *in vitro* ubiquitination assays with purified UPP components (E1, E2, ubiquitin, and ATP) along with EpE89-uAb as the E3 enzyme and ERK2 as the target (note that ERK2 has 23 lysine residues in addition to its N-terminus that serve as potential ubiquitin attachment sites) (**Fig. 2b**). UbcH5α was used as the E2 enzyme because it has previously been shown to function with CHIP *in vitro* ^22, 45^. High-molecular-weight (HMW) bands corresponding to ubiquitinated ERK2 were detected with the pan-ERK antibody, which correlated with the appearance of HMW ubiquitin species that were detected with the anti-ubiquitin antibody (**Fig. 2b**). The intensity of the HMW bands became more pronounced at later incubation times and was characteristic of CHIP-mediated polyubiquitination of its natural and unnatural targets ^22, 45^. Similar ubiquitination results were observed for pE59-uAb and E40-uAb (**Supplementary Fig. 2c**). Only when all UPP components were included in the reaction was ubiquitination observed and neither the binding domain, unfused EpE89, nor the catalytic domain, CHIPΔTPR, was capable of producing ubiquitinated ERK2 (**Fig. 2c**). Collectively, these results confirm that the CHIPΔTPR domain retained E3 ligase activity in the context of the uAb chimeras and was capable of directly transferring ubiquitin to ERK2.

### Ubiquibodies efficiently degrade exogenous and endogenous ERK

To characterize the degradation potential of pan-ERK-specific EpE89-uAb, we first evaluated soluble expression in mammalian cells. Specifically, wild-type (wt) HEK293T cells were transiently transfected with plasmid DNA encoding the chimeric EpE89-uAb construct and cell lysate was prepared 24 h post-transfection. Strong expression of EpE89-uAb was detected in soluble lysates by immunoblot analysis using an anti-His6x antibody (**Supplementary Fig. 3a**). Interestingly, while pE59-uAb also exhibited strong soluble expression, the E40-uAb construct was barely detectable. To determine whether this poor expression was somehow related to the cell line, we also expressed the uAbs in MCF7 breast cancer cells and observed an identical expression pattern (**Supplementary Fig. 3b**). In light of these poor steady-state levels observed for E40-uAb, we focused our attention on the EpE89-uAb and pE59-uAb constructs hereafter. It is also worth mentioning that expression of the three unfused DARPins was barely detectable under the conditions tested, providing additional evidence for the ability of the CHIPΔTPR domain to enhance soluble expression and revealing an unexpected benefit arising from uAb chimeragenesis.

To investigate intracellular knockdown, we next leveraged an exogenously expressed ERK2-EGFP reporter fusion. Specifically, a previously engineered cell line that stably expresses an ERK2-EGFP-encoding transgene (HEK293T^ERK2-EGFP^) ^27^, were transiently transfected with plasmid DNA encoding the uAbs. Immunoblot analysis of lysate derived from these cells revealed that expression of both EpE89-uAb and pE59-uAb promoted efficient clearance of ERK2-EGFP relative to the steady-state level observed in the same cells transfected with an empty plasmid or plasmid DNA encoding either CHIPΔTPR or Off7-uAb (**Supplementary Fig. 3c**). The depletion of ERK2-EGFP protein levels by EpE89-uAb and pE59-uAb coincided with an overall reduction of GFP fluorescence as determined by flow cytometric analysis (**Supplementary Fig. 3c**). The extent of fluorescence reduction associated with ERK2-GFP knockdown was reminiscent of that observed previously for EGFP-HRAS, EGFP-KRAS, and SHP2-EGFP using uAbs comprised of synthetic binding proteins against HRAS, KRAS and SHP2, respectively ^27^.

While the above results demonstrated the feasibility for uAb-mediated knockdown of an ERK2-containing fusion protein in living cells, we cannot rule out the possibility that ubiquitin was conjugated exclusively to the EGFP domain and not on ERK1/2, which would limit the practical utility of EpE89-uAb and pE59-uAb for proteolytic silencing of untagged ERK forms. Interestingly, the pan-specific ERK antibody used to detect ERK2-EGFP also revealed depletion of endogenous ERK proteins in the lysates derived from cells expressing EpE89-uAb and pE59-uAb (**Supplementary Fig. 3c**), suggesting that the uAbs could indeed accelerate the turnover of unmodified ERK in addition to its EGFP-tagged counterpart. However, even this result was inconclusive as endogenous ERK1 and ERK2 are known to homodimerize ^46^, which leaves open the possibility that endogenous ERK2 could heteroassemble with ubiquitinated ERK2-EGFP (where again the ubiquitin might be installed on EGFP only) and become targeted for proteolysis via a piggy-back mechanism.

Therefore, we focused our attention on determining the extent to which endogenously expressed, unmodified ERK could be degraded by EpE89-uAb and pE59-uAb. To this end, wt HEK293T cells were transiently transfected with the EpE89-uAb and pE59-uAb-encoding plasmids. After 24 h, transfected cells displayed dramatically reduced steady-state levels of total ERK1/2 compared to cells receiving empty plasmid DNA as detected by the phosphorylation-insensitive pan-ERK antibody (**Fig. 3a and b**). Because cytoplasmic ERK is present as a mixture of nonphosphorylated and phosphorylated forms in both non-stimulated and stimulated HEK293T cells ^39, 47^, we also probed lysates with an anti-pERK1/2 antibody. Consistent with their strong pERK2 binding activity, both uAbs showed potent reduction of pERK levels that mirrored total ERK knockdown, with pE59-uAb promoting greater reduction of pERK (**Supplementary Fig. 4**). Importantly, transfection of HEK293T cells with either unfused EpE89, pE59, or CHIPΔTPR resulted in little to no change in total ERK1/2 protein levels, indicating that none of these domains alone was capable of target depletion and confirming the importance of the bifunctional uAb design. The non-specific Off7-uAb was also incapable of promoting ERK1/2 degradation, thereby validating the targeted nature of ERK depletion by EpE89-uAb and pE59-uAb. As was seen above, soluble expression of the DARPins was greatly enhanced by fusion to CHIPΔTPR. It is also noteworthy that the levels of a housekeeping protein, β-tubulin, and the native CHIP substrate, Hsp70, were not affected by expression of EpE89-uAb, pE59-uAb, or any of the other control constructs.

**Figure 3.**
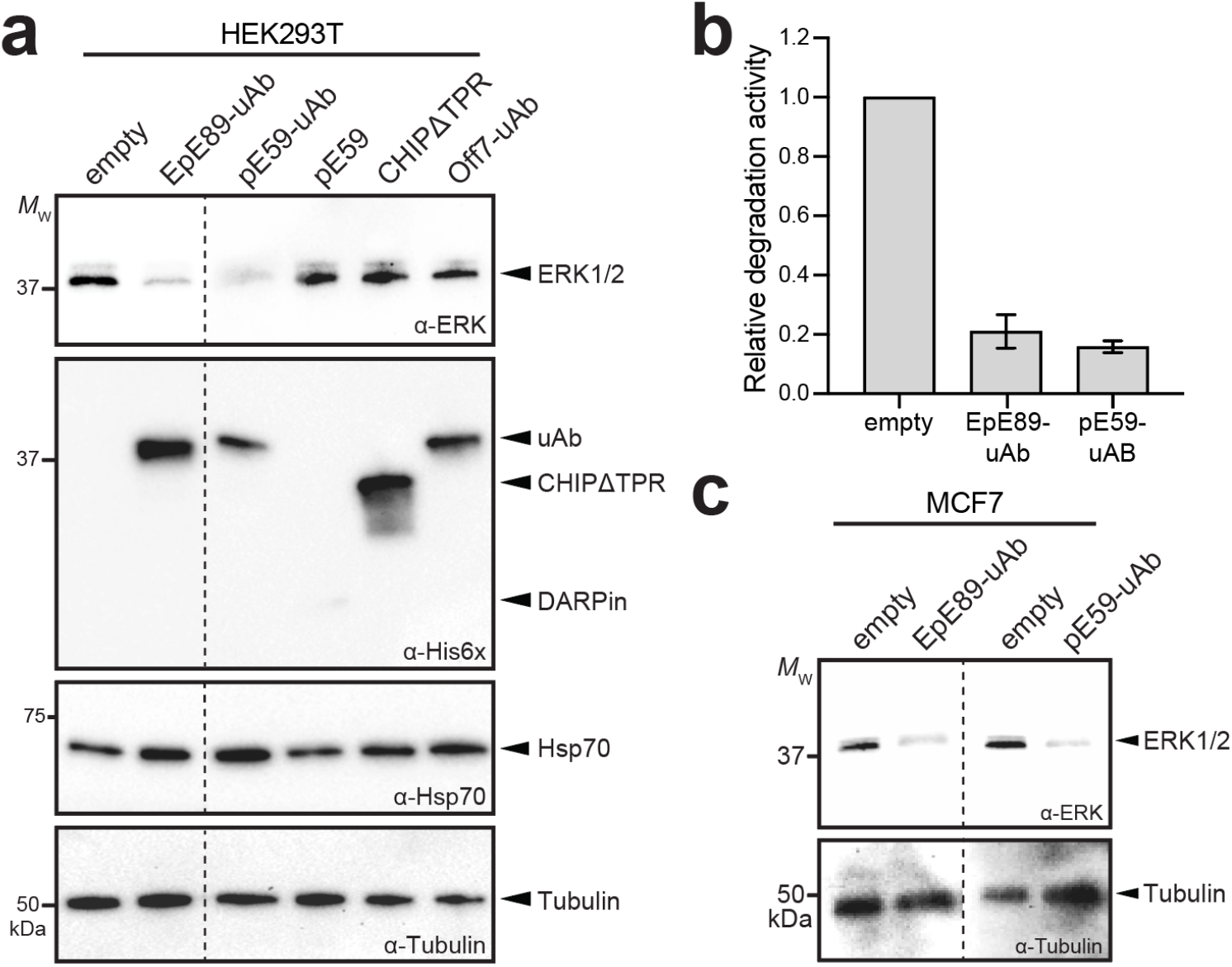
Engineered uAbs efficiently degrade endogenous ERK in living cells. (a) Immunoblot analysis of extracts prepared from HEK293T cells transfected with empty pcDNA3 or pcDNA3 encoding each of the constructs indicated at 0.25 μg plasmid DNA per well. Cells were harvested 24 h post-transfection, after which extracts were prepared and subjected to immunoblotting. Blots were probed with the following: pan-ERK antibody (α-ERK) to detect total ERK1/2 expression; polyhistidine antibody (α-His6x) to detect uAbs, DARPins, and CHIPΔTPR constructs; and Hsp70-specific antibody (α-Hsp70) to detect native CHIP substrate. Lanes were normalized by total protein content and equivalent loading was confirmed by probing with β-tubulin (α-Tub). Molecular weight (*M*_W_) markers are indicated at left. Results are representative of at least three biological replicates. (b) Relative quantitation of total ERK1/2 levels by densitometry analysis of α-ERK immunoblot images using ImageJ Software. Intensity data for uAb bands was normalized to band intensity for empty plasmid control cases from six independent experiments. Error bars represent standard deviation of the mean. (c) Immunoblot analysis of extracts prepared from MCF7 cells transfected with empty pcDNA3 or pcDNA3 encoding EpE89-uAb or pE59-uAb at 0.25 μg plasmid DNA per well. Blots prepared as in (a). Dashed line indicates splicing of the same blot.

We next explored whether our anti-ERK approach would work in other cell lines. Specifically, we investigated the ability of EpE89-uAb and pE59-uAb to degrade ERK in MCF7 breast cancer cells, which have served as a useful model for studying ERK expression, activation and signaling ^48, 49^. Akin to the results with HEK293T, we observed strong reduction of total ERK1/2 levels in MCF7 cells transfected with plasmid DNA encoding the EpE89-uAb and pE59-uAb constructs compared to cells transfected with empty plasmid (**Fig. 3c**). Degradation was most pronounced at 24 h post-transfection; however, clearly visible depletion of ERK1/2 persisted out to 48 and 72 h (**Supplementary Fig. 3b**), consistent with the duration of uAb-mediated GFP silencing observed in our previous work ^27^.

### Pan-specific uAbs transfer ubiquitin to distinct sites on ERK and pERK

To further elucidate the origins of pan-specific degradation, we profiled the ubiquitination patterns generated by EpE89-uAb and pE59-uAb on nonphosphorylated and phosphorylated ERK2. Specifically, *in vitro* ubiquitination reactions were performed with each of the uAbs in the presence of either ERK2 or pERK2 as substrates, after which HMW products (~50-250 kDa) were separated by SDS-PAGE, excised from the gel, digested with trypsin, and analyzed by liquid chromatography-tandem mass spectrometry (LC-MS/MS; **Fig. 4a**). Trypsin digestion of a ubiquitinated protein leaves the C-terminal glycine-glycine residues of ubiquitin attached to the ubiquitinated lysine residue ^50^. Therefore, we searched the MS data for this modification on ERK2/pERK2 peptides and identified Lys residues to which ubiquitin was conjugated. In general, the ubiquitination profiles of EpE89-uAb and pE59-uAb were highly similar, with both uAbs transferring ubiquitin to multiple lysine residues in ERK2 and pERK2 (**Fig. 4b**). These overlapping profiles help to explain the observed pan-specific ERK degradation of each uAb. Moreover, both uAbs preferentially ubiquitinated one face of ERK2/pERK2 (oriented forward in **Fig. 4c**), consisting of the plane formed by the N- and C-lobes near the active site of ERK2. This face was aligned with the positioning of the C-terminus of the DARPins with ERK2 as seen in co-crystal structures ^39^ and would thus be located in closest proximity to CHIPΔTPR when bound by the uAb chimera (**Fig. 1b**).

**Figure 4.**
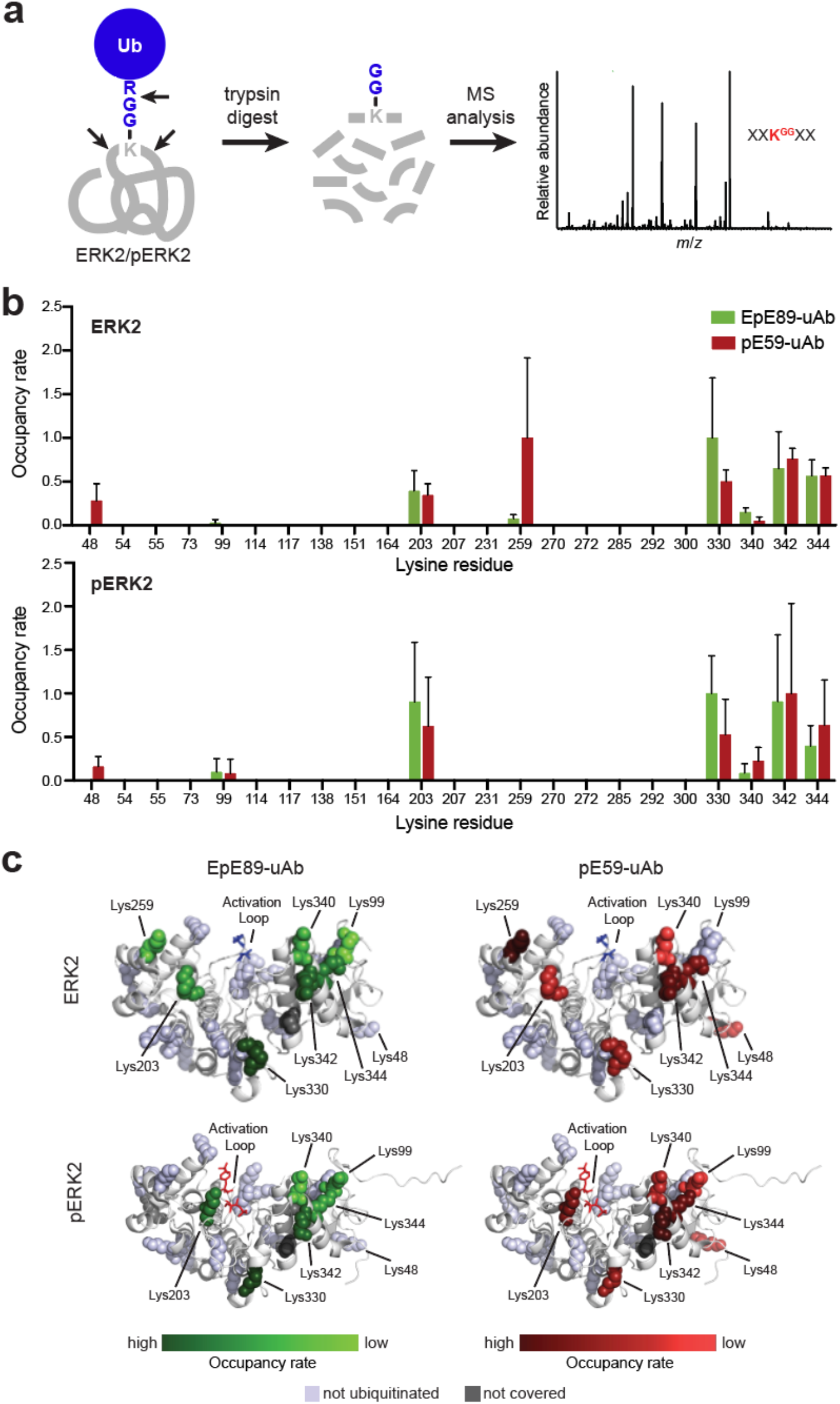
Engineered uAbs install ubiquitin on multiple lysine residues in ERK2 and pERK2. (a) Schematic of ubiquitin profiling experiment for revealing precise ubiquitination sites in ERK2/pERK2. Briefly, mass spectrometry can be used to identify ubiquitin attachment sites based on the characteristic mass shift caused by the presence of diglycine (GG) that is retained on ubiquitinated lysine residues within peptides after trypsin digestion. (b) Occupancy rate of GG modification of ERK2/pERK2 lysine residues by EpE89-uAb and pE59-uAb by LC-MS/MS. Peptides corresponding to 80% of the ERK2/pERK2 sequences were identified using Mascot software. Data were generated by normalizing ubiquitinated residue counts relative to total residue counts, and by averaging across three independent experiments. Ubiquitinated peptide counts of peptides containing more than one non-C-terminal lysine residue were averaged over all non-C-terminal lysines. (c) Mapping of ubiquitination sites on ERK2 and pERK2 where ERK2/pERK2 backbones are shown as white ribbons and ubiquitinated lysines represented as spheres colored by heat map as indicated. Lysines not covered by mass spectrometry analysis (dark grey spheres) and lysines not identified as ubiquitinated (light grey spheres) are also depicted. Structures adapted from PDB ID: 3ZU7 (ERK2) and PDB ID: 3ZUV (pERK2) of Kummer *et al*. ^39^ using PyMOL software.

Of the 23 total lysines in ERK2, 7 sites (K48, K99, K203, K330, K340, K342, and K344) were found to be modified in both ERK2 and pERK2 (**Fig. 4b and c**). Only one additional residue, K259, was ubiquitinated in ERK2 and not pERK2, suggesting that the conformational change upon phosphorylation may reposition K259 away from the U-box-bound, ubiquitin-charged E2, UbcH5α. This site was also interesting because it was much more frequently modified by pE59-uAb than EpE89-uAb in ERK2 and was not modified by either uAb in pERK2. Four of the modified lysine residues (K99, K340, K342, and K344) were clustered in the three-dimensional structure of ERK2 (**Fig. 4c**), providing clues about the orientation of the charged E2-uAb complex relative to the target surface and consistent with the predicted plane of ubiquitination (**Fig. 1b**). Residues K203 and K330 were among the most frequently ubiquitinated despite being positioned away from the plane of ubiquitination, suggesting mobility of the E2-uAb complex as has been observed previously for native E2-E3 complexes ^51^. Residue K48 was one of the least often ubiquitinated sites and the only lysine not on the same face of ERK to be ubiquitinated. Interestingly, K48 in both ERK2 and pERK2 was modified by EpE89-uAb but not at all by pE59-uAb, suggesting that the DARPin domains recognize different epitopes and thus differentially orient the uAbs with respect to ERK2/pERK2 in a manner that affects how the substrate is ubiquitinated.

The polyubiquitin chain topology formed by the uAbs was analyzed using an identical LC-MS/MS approach (**Supplementary Fig. 5a**), which identified the isopeptide linkages between the terminal carboxyl group of a free ubiquitin molecule and one of seven lysine residues present in a substrate-attached ubiquitin. According to this analysis, both uAbs produced nearly identical ubiquitin chain topologies, preferentially forming K6, K11, K48, and K63 polyubiquitin linkages in the presence of the E2 UbcH5α (**Supplementary Fig. 5b**). These same linkages were observed previously on natural and unnatural substrates that had been ubiquitinated *in vitro* by full-length CHIP and CHIP-based uAbs, respectively ^22, 52^. These results are significant from a targeted degradation standpoint, as K48 serves as the principal recognition signal for the 26S proteasome and generally induces substrate degradation ^7^, while K6, K11 and K63 have also been implicated as proteasomal targeting signals ^53, 54^. Taken together, these results provide clear evidence for highly similar ERK2/pERK2 ubiquitination by EpE89-uAb and pE59-uAb, thereby providing a convenient explanation for their comparable pan-specific ERK degradation.

## Discussion

Ubiquibodies are a customizable proteome editing technology for inducing targeted proteolysis of intracellular proteins and thus hold great potential as both a research tool for dissecting protein networks and as a therapeutic modality with the potential for inhibiting drug targets that have so far evaded pharmacological intervention. In this study, we engineered chimeric uAbs comprised of the human E3 ubiquitin ligase CHIP, and different ERK-specific DARPins that were capable of accelerating the turnover of exogenous or endogenous ERK protein kinase. In particular, two of the uAbs were shown to be global ERK degraders, redirecting all ERK1/2 proteoforms, including both active (doubly phosphorylated) and inactive (nonphosphorylated) conformations, to the 26S proteasome for degradation in different cell lines including MCF7 breast cancer cells. These results add to a growing body of evidence that reveals the effectiveness of designer uAb constructs in promoting the clearance of POIs ^22, 26–34^ including some that have been classified as difficult-to-drug.

As we demonstrated here and in previous works, the combination of synthetic binding proteins having affinity and specificity for the POI with the catalytic domain of E3 ligases opens the door to targeted knockout of intracellular proteins and their posttranslationally modified isoforms. Indeed, numerous structurally diverse POIs that span a broad range of molecular weights (from 27*-*179 kDa) and subcellular locations (i.e., cytoplasm, nucleus, membrane-associated, and transmembrane) have been targeted for degradation using uAb technology ^18–22, 27^. Importantly, design and construction of uAbs does not require knowledge of the biological function or interaction partners of the POI. Instead, uAbs take advantage of synthetic binding proteins that have already been developed or emerge anew such as from systematic, genome-wide efforts to generate and validate *de novo* protein binders against the human proteome ^25^. Because obtaining antibody mimetics that bind with high specificity and affinity to a target is generally easier than obtaining small molecules with the same properties, making custom-designed uAbs from scratch should be more straightforward than generating new PROTACs ^3, 55^.

The depletion of total ERK pools obtained with EpE89-uAb was expected given its affinity for both ERK and pERK; however, the ability of pE59-uAb to also function as a pan-specific degrader was somewhat surprising given its reported specificity for pERK ^39^. We suspect that despite its clear preference for pERK, pE59-uAb may bind non-cognate ERK2 with enough affinity to still promote efficient substrate turnover. Indeed, the unfused pE59 DARPin is known to bind non-cognate ERK2 with micromolar affinity (*K*_D_ = 3.5–8.7 μM ^39^), which should be sufficient to promote ubiquitin transfer given that the measured affinity between CHIP and its native substrates Hsp70, Hsp90, and Hsc70 is also in the low micromolar range (*K*_D_ = 0.3–2.3 μM) ^56^. Moreover, the binding of ERK2 by pE59-uAb is likely to be enhanced by avidity effects that arise from dimerization of the CHIP-based uAb. Importantly, while pan-specific degraders were generated here that promoted degradation of multiple proteoforms, uAbs have also been created that selectively degrade distinct forms of a protein ^17, 18, 27^. Collectively, the designer binding of uAbs could open up new avenues for disease intervention by ablating either the entire family of functionally overlapping proteins or a specific posttranslational event that is preferentially dysregulated in a diseased state. In the case of ERK, it has been proposed that ERK1/2-selective inhibitors could provide potential therapeutic opportunities for a broad spectrum of cancers bearing RAS, RAF, and MEK mutations ^36, 57^. Given the functional redundancy of ERK1 and ERK2, broad inactivation of both family members may be needed to inhibit cellular proliferation and causes apoptosis in tumor cells and induce significant tumor regression, a hypothesis that could be investigated using our pan-ERK-specific degraders. To this end, it should be pointed out that promising *in vivo* results have been obtained using experimental viral and non-viral vectors to deliver uAb genes ^20, 27, 58, 59^, indicating that clinical translation may not be that far off.

## Material and Methods

### Plasmid construction

*E. coli* strain DH5α was used for the construction and propagation of all plasmids. The creation of plasmids encoding uAb constructs and related controls were generated following published protocols ^34^. Genes encoding each of the DARPins were PCR amplified from pDST67-based plasmids encoding EpE89, pE59, and E40 ^39^ using primers that introduced NcoI and EcoRI overhangs. The resulting PCR amplicons were ligated in plasmid pET28a-R4-uAb ^22^, which had been doubly digested with NcoI/EcoRI to excise the gene encoding scFv13-R4 (R4). This process yielded plasmids pET28a-EpE89-uAb, pET28a-pE59-uAb, and pET28a-E40-uAb, which encoded each of the DARPins followed by a flexible GSGSG linker and then CHIPΔTPR bearing a tandem FLAG-His6x sequence at its C-terminus. A similar strategy was used to generate plasmid pET28a-Off7-uAb, where the gene encoding Off7 was PCR amplified from plasmid pRH-DsbA-off7 ^60^ (kind gift from Mark Ostermeier, Johns Hopkins University). To generate plasmid pET28a-CHIPΔTPR for expression of unfused CHIPΔTPR, a gene fragment corresponding to amino acids 128-303 of human CHIP was PCR amplified with primers that introduced NcoI and SalI overhangs and ligated into the same sites in plasmid pET28a-R4-uAb that had been doubly digested with NcoI/SalI to excise the R4-uAb while leaving behind the tandem FLAG-His6x sequence. To generate plasmids for expression of unfused DARPins, genes encoding each of the DARPins were similarly PCR amplified from pDST67-based plasmids using primers that introduced NcoI and HindIII overhangs as well as an N-terminal RGS-His6x sequence. The resulting PCR amplicons were cloned into pET28a(+) between NcoI and HindIII, yielding plasmids pET28a-EpE89, pET28a-pE59, and pET28a-E40. For expression in human cell lines, all uAbs and control proteins were cloned into plasmid pcDNA3, a mammalian expression vector with constitutive CMV promoter. This involved PCR amplification of the target genes using the respective pET28a-based vectors described above as template along with primers that introduced HindIII and XbaI overhangs and a Kozak sequence at the start codon. The resulting PCR amplicons were then ligated between the HindIII and XbaI sites in pcDNA3 to yield the desired plasmids including pcDNA3-EpE89-uAb, pcDNA3-pE59-uAb, and pcDNA3-E40-uAb. All plasmids were confirmed by DNA sequencing at the Biotechnology Resource Center (BRC) Genomics Facility at the Cornell Institute of Biotechnology

### Protein expression and purification

All purified uAbs, unfused DARPins, and CHIPΔTPR were obtained from cultures of *E. coli* BL21(DE3) cells carrying pET28a-based vectors grown in Luria-Bertani (LB) medium. Expression was induced with 0.1 mM IPTG when the culture density (Abs_600_) reached 0.6-0.8 and proceeded for 6 hr at 30 °C, after which cells were harvested by centrifugation at 4,000×g for 20 min at 4 °C. The resulting pellets were stored at –80 °C overnight. Thawed pellets were resuspended in 15 mL phosphate-buffered saline (PBS) supplemented with 10 mM imidazole (pH 7.4) and lysed with a high-pressure homogenizer (Avestin EmulsiFlex-C5). Lysates were cleared of insoluble material by centrifugation at 20,000×g for 20 min at 4 °C. Clarified lysates containing His6x-tagged proteins were subjected to gravity-flow Ni^2+^-affinity purification using HisPur Ni-NTA Resin (ThermoFisher) following manufacturer’s protocols. Elution fractions were desalted into PBS buffer (pH 7.4) using PD-10 Desalting Columns (Cytiva) following manufacturer’s protocols. Purified proteins were stored at 4 °C for up to two weeks or diluted to 25% (v/v) glycerol and stored indefinitely at −80 °C. Final purity of all proteins was confirmed by SDS-polyacrylamide gel electrophoresis (PAGE) and Coomassie staining. Purity of all proteins was typically >95%.

Purified uAbs and CHIPΔTPR were subjected to SEC analysis as described previously ^61^. Standards used to calibrate the SEC column were a lyophilized mix of thyroglobulin, bovine γ-globulin, chicken ovalbumin, equine myoglobin, and vitamin B12, MW 1,350–670,000, pI 4.5–6.9 (BioRad). Proteins were stored at a final concentration of 1 mg/mL in SEC buffer (20 mM Tris pH 7.5, 50 mM NaCl, 1 mM EDTA pH 8.0) at 4 °C.

To produce biotinylated ERK2 and pERK2 proteins, strain BL21(DE3) was co-transformed with plasmid pSPI03-BirA-His ^62^ (kind gift from Amy Karlsson, University of Maryland) along with either plasmid pLV-ERK2-Avi for expressing nonphosphorylated ERK2 or pLV-MEK1R4F-ERK2-His-Avi for expressing doubly phosphorylated ERK2, respectively ^39^. These latter plasmids introduced N-terminal Avi tags on ERK2 and pERK2 for biotinylation *in vivo* by the biotin ligase BirA encoded in plasmid pSPI03-BirA-His and C-terminal His6x tags for affinity purification and immunodetection. Following expression, bacterial cell pellets were harvested by centrifugation, pelleted, and resuspended in PBS (pH 7.4) with 1 mM DTT and 0.05% Tween-20. The resulting cell suspensions were homogenized as above, after which the clarified lysates containing biotinylated ERK2 and pERK2 were subjected to avidin agarose (ThermoFisher) to purify the Avi-tagged proteins according to manufacturer’s protocols. Following elution with 2 mM biotin, the eluents were subjected to Ni^2+^-affinity purification as above to remove free biotin and further enhance the purity. Biotinylated ERK2 and pERK2 were analyzed by SDS-PAGE followed by Coomassie staining to confirm purity, which was typically >95% for both proteins.

### Affinity precipitation

Affinity purification was performed as described ^63^. Briefly, purified uAbs, unfused DARPins, and CHIPΔTPR were captured on HisPur Ni-NTA Resin (ThermoFisher) by incubating 300 μg of each protein with 1-mL resin slurry for 30 min at 4 °C with end-over-end rotation. Prepared resin was incubated with 10 μL of lysate at 4 °C overnight. Resin was washed with PBS supplemented with 25 mM imidazole (pH 7.4), and proteins were eluted with PBS supplemented with 250 mM imidazole (pH7.4). Samples were boiled with 2× Laemmli loading buffer and analyzed by immunoblotting as described below.

### Protein analysis

Proteins were separated using Precise Tris-HEPES 4–20% SDS-polyacrylamide gels (ThermoFisher). Coomassie R-250 stain (BioRad) was used to visualize proteins in SDS-PAGE. Immunoblotting was performed according to standard protocols. Following transfer of proteins, polyvinylidene fluoride (PVDF) membranes were probed with the following antibodies at 1/2500 or 1/5000 dilution: rabbit anti-p44/42 MAPK (ERK1/2) antibody (Cell Signaling, cat # 4695 S) to detect ERK2; rabbit anti-p-p44/42 MAPK (ERK1/2) (Cell Signaling, cat # 9101 S) to detect pERK2; mouse anti-ubiquitin (Millipore, cat # P4D1-A11) to detect ubiquitin; rabbit anti-Lys27 (Abcam, cat # ab238442) to detect K27-linked ubiquitin; rabbit anti-Lys48 (Millipore, cat # Apu2) to detect K48-linked ubiquitin; rabbit anti-Lys63 (Millipore, cat # Apu3) to detect K48-linked ubiquitin; mouse anti-Hsp70 (Enzo Life Sciences, cat # C92F3A) to detect Hsp70; rabbit anti-β-tubulin (Cell Signaling Technology, cat # 5346) to detect β-tubulin; rabbit anti-FLAG-HRP (Abcam, cat # ab49763) to detect uAbs and CHIPΔTPR; and rabbit anti-His6-HRP (Abcam; cat # ab1187) to detect uAbs, unfused DARPins, and CHIPΔTPR.

### ELISA

To analyze binding to purified ERK and pERK, ELISA analysis was performed as described previously ^39^. Briefly, biotinylated ERK2 and pERK2 (100 nM) were immobilized on NeutrAvidin-coated 96-well plates (Pierce) overnight at 4 °C and then washed twice with PBS (pH 7.4) supplemented with 1 mM DTT and 0.05%Tween-20. Next, the plates were blocked for 1 h with PBS (pH 7.4) supplemented with 1 mM DTT, 0.05% Tween-20, and 1% (w/v) BSA. All subsequent ELISA steps were performed at 4 °C in PBS (pH 7.4) with 1 mM DTT and 0.05% Tween-20. To measure binding activity, varying concentrations of purified uAbs, unfused DARPins, or CHIPΔTPR were applied wells with or without ERK2 or pERK2 for 1 h. Following three washes, binding activity was detected by rabbit anti-His6-HRP (Abcam; cat # ab1187) or mouse anti-RGS-His antibody (Qiagen; cat # 34610) at 1:5000 dilution followed by goat anti-rabbit-HRP conjugate (Abcam; ab6789) at 1:2500 dilution. After 1 h of incubation at room temperature, plates were washed and then incubated with SigmaFast OPD HRP substrate (Sigma) for 30 min in the dark. The reaction was quenched with 3 M H_2_SO_4_ and the absorbance of the wells measured at 492 nm.

### Ubiquitination assays

Ubiquitination assays were performed as previously described ^45^ in the presence of 0.1 μM purified human UBE1 (Boston Biochem), 4 μM human UbcH5α/UBE2D1 (Boston Biochem), 3 μM uAb (or equivalent control protein), 1.5 μM human ERK2 or phosphoERK2 (ProQinase), 50 μM human ubiquitin (Boston Biochem), 4 mM ATP and 1 mM DTT in 20 mM MOPs, 100 mM KCl, 5 mM MgCl_2_, pH 7.2. Reactions were carried out at 37 °C for 2 h (unless otherwise noted) and stopped by boiling in 2× Laemmli loading buffer for analysis by immunoblotting.

### Flow cytometric analysis

Cells were passed into 12-well plates at 10,000 cells/cm^2^. At 16-24 h after seeding, cells were transiently transfected as described above. Culture media was replaced 4–6 h post-transfection. Then, 24 h post-transfection, cells were harvested and resuspended in PBS for analysis using a FACSCalibur (BD Biosciences). FlowJo software (Version 10) was used to analyze samples by geometric mean fluorescence determined from 10,000 events.

### Cell culture, transfection, and lysate preparation

HEK293T and MCF7 cell lines were obtained from ATCC, while the HEK293T^ERK2-EGFP^ cell line was previously generated in-house ^27^. HEK293T and HEK293T^ERK2-EGFP^ cells were cultured in DMEM media supplemented with high glucose and L-glutamine (VWR) supplemented with 10% Hyclone FetalClone I serum (VWR) and 1% penicillin-streptomycin-amphotericin B (ThermoFisher). MCF7 cells were cultured similarly but insulin (10 mg/mL, Sigma) was added to the media. All cells were maintained at 37 °C, 5% CO_2_ and 90% relative humidity (RH). Additionally, all cell lines were maintained at low passage numbers and routinely checked for *Mycoplasma* by PCR according to standard procedures. Cells were transfected in 6-well dishes at 60-80% confluency with 2 μg total plasmid DNA using empty pcDNA3 plasmid to balance all transfections. Transfection was performed using jetPRIME^®^(Polyplus Transfection) according to manufacturer’s instructions with a 1:2 ratio (w/v) of jetPRIME^®^ to DNA with growth media refreshed at 4 h post-transfection. At 24 h post-transfection, cell lysate was prepared by harvesting cells in PBS, pelleting at 8000×g for 5 min at 4 °C, and freezing at −20 °C until analyzed by immunoblotting. Thawed pellets were lysed in NP40 lysis buffer (150 mM NaCl, 1% Nonidet P-40, 50 mM Tris-HCl, pH 7.4) by pipetting and mixing for 30 min at 4 °C. Soluble fractions were obtained by centrifugation of lysed cells at 18,000×g for 20 min at 4 °C. Samples were boiled in 2× Laemmli sample buffer for analysis by immunoblotting.

### Mass spectrometry analysis

For LC-MS/MS sample preparation, ubiquitination assays were performed as described above. Reactions were resolved by SDS-PAGE and stained by Coomassie R250 prior to gel excision. The protein bands were excised from an SDS-PAGE gel, cut into ~1-mm^3^ cubes, and submitted to the Biotechnology Resource Center (BRC) Proteomics and Metabolomics Facility at the Cornell Institute of Biotechnology for further analysis. Specifically, the gel bands were washed in 200 μL of deionized water for 5 min, followed by 200 μL of 100 mM ammonium bicarbonate/acetonitrile (1:1) for 10 min, and finally 200 μL of acetonitrile for 5 min. The acetonitrile was discarded, and the gel bands were dried in a speed-vac for 10 min. The gel pieces were rehydrated with 70 μL of 10 mM DTT in 100 mM ammonium bicarbonate and incubated for 1 h at 56 °C. The samples were allowed to cool to room temperature, after which 100 μL of 55 mM iodoacetamide in 100 mM ammonium bicarbonate was added and the samples were incubated at room temperature in the dark for 60 min. Following incubation, the gel slices were again washed as described above. The gel slices were dried and rehydrated with 50 μL of trypsin at 50 ng/μL in 45 mM ammonium bicarbonate and 10% acetonitrile on ice for 30 min. The gel pieces were covered with an additional 25 μL of 45 mM ammonium bicarbonate and 10% acetonitrile, and incubated at 37 °C for 19 h. The digested peptides were extracted twice with 70 μL of 50% acetonitrile, 5% formic acid (vortexed 30 min and sonicated 10 min) and once with 70 μL of 90% acetonitrile, 5% formic acid. Extracts from each sample were combined and lyophilized.

The lyophilized in-gel tryptic digest samples were reconstituted in 20 μL of nanopure water with 0.5% formic acid for nanoLC-ESI-MS/MS analysis, which was carried out by a LTQOrbitrap Velos mass spectrometer (ThermoFisher) equipped with a CorConneX nano ion source device (CorSolutions LLC). The Orbitrap was interfaced with a nano HPLC carried out by an UltiMate3000 UPLC system (Dionex). The gel extracted peptide samples (2–4 μL) were injected onto a PepMap C18 trap column-nano Viper (5 μm, 100 μm × 2 cm, Thermo Dionex) at 20 μL/min flow rate for online desalting and then separated on a PepMap C18 RP nanocolumn (3 μm, 75 μm × 15 cm, Thermo Dionex) which was installed in the “Plug and Play” device with a 10-μm spray emitter (NewObjective). The peptides were then eluted with a 90-min gradient of 5% to 38% acetonitrile in 0.1% formic acid at a flow rate of 300 nl/min. The Orbitrap Velos was operated in positive ion mode with nanospray voltage set at 1.5 kV and source temperature at 275 °C. Internal calibration was performed with the background ion signal at *m/z* 445.120025 as the lock mass. The instrument was operated in parallel data-dependent acquisition mode using FT mass analyzer for one survey MS scan for precursor ions followed by MS/MS scans on top 7 highest intensity peaks with multiple charged ions above a threshold ion count of 7,500 in both LTQ mass analyzer and high-energy collision dissociation (HCD)-based FT mass analyzer at 7,500 resolution. Dynamic exclusion parameters were set at repeat count 1 with a 15-s repeat duration, exclusion list size of 500, 30-s exclusion duration, and ±10 ppm exclusion mass width.

HCD parameters were set at the following values: isolation width of 2.0 *m*/*z*, normalized collision energy of 35%, activation *Q* at 0.25, and activation time of 0.1 ms. All data were acquired using Xcalibur operation software (version 2.1, ThermoFisher).

All MS and MS/MS raw spectra were processed and searched using Proteome Discoverer 1.3 (PD1.3; ThermoFisher) against databases downloaded from the NCBI database. The database search was performed with two-missed cleavage site by trypsin allowed. The peptide tolerance was set to 10 ppm, and MS/MS tolerance was set to 0.8 Da for collision-induced dissociation and 0.05 Da for HCD. A fixed carbamidomethyl modification of cysteine, variable modifications on methionine oxidation, and ubiquitin modification of lysine were set. The peptides with low confidence score (with an Xcorr score < 2 for doubly charged ion and < 2.7 for triply charged ion) defined by PD1.3 were filtered out, and the remaining peptides were considered for the peptide identification with possible ubiquitination determinations. All MS/MS spectra for possibly identified ubiquitination peptides from initial database searching were manually inspected and validated using both PD1.3 and Xcalibur (version 2.1) software.

## Data availability

All data generated or analyzed during this study are included in this article (and its supplementary information) or are available from the corresponding authors on reasonable request.

## Acknowledgements

We thank Dr. Cam Patterson, Dr. Mark Ostermeier, Dr. Amy Karlsson, and Dr. Melanie Cobb for plasmids used in this study. We also thank Dr. Peter Schweitzer and the BRC Genomics Facility at the Cornell Institute of Biotechnology for sequencing experiments and Sheng Zhang and the Proteomics and Metabolomics Facility of the Biotechnology Resource Center of Cornell Institute of Biotechnology for help with mass spectrometry experiments. This work was supported by the National Science Foundation Grant CBET-1605242 (to M.P.D.), the National Institutes of Health Grant Numbers R21CA132223 and R01GM137314 (to M.P.D.), the Defense Threat Reduction Agency HDTRA1-20-10004 (to M.P.D.), the New York State Office of Science, Technology and Academic Research Distinguished Faculty Award (to M.P.D.), and the Cornell Technology Acceleration and Maturation (CTAM) Fund. The work was also supported by seed project funding (to M.P.D.) through the National Institutes of Health-funded Cornell Center on the Physics of Cancer Metabolism (supporting grant 1U54CA210184-01). The content is solely the responsibility of the authors and does not necessarily represent the official views of the National Cancer Institute or the National Institutes of Health. E.A.S. and M.B.L. were each supported by National Science Foundation Graduate Research Fellowships (grants DGE-1650441 and DGE-1144153, respectively) and Cornell Presidential Life Science Fellowships. M.B.L. was also supported by a Cornell Fleming Graduate Scholarship. B.M. was supported by a Royal Thai Government Fellowship.

## Author Contributions

E.A.S. designed research, performed all research, analyzed all data and wrote the paper. M.B.L., B.M., M.L., and K.J.F. performed research. T.Y. and C.M. helped write and edit the paper. L.K., and A.P. aided in data interpretation. M.P.D. directed research, analyzed data and wrote the paper.

