## Supplementary Information for "Engineering single pan-specific ubiquibodies for targeted degradation of all forms of endogenous ERK protein kinase"

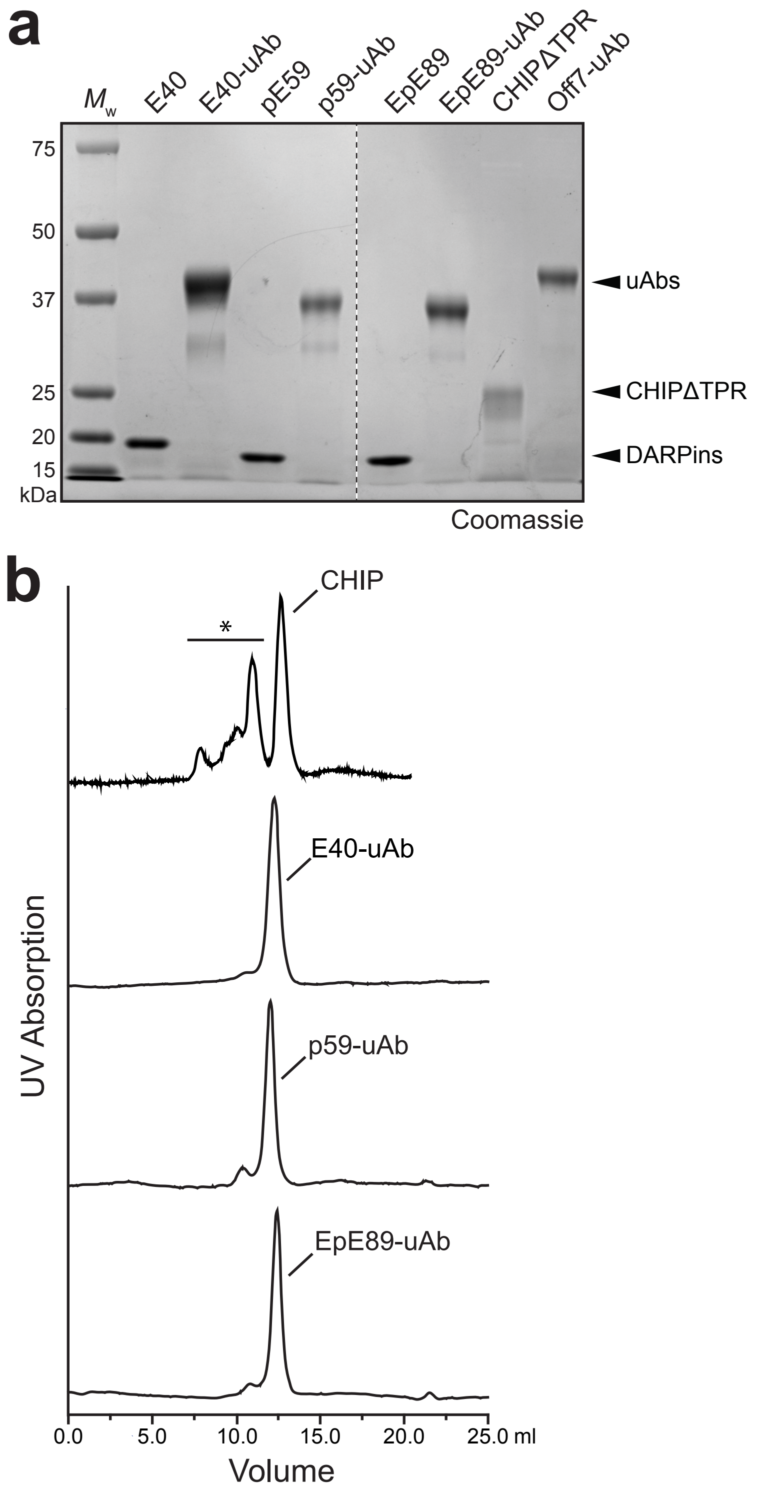


**Supplementary Figure 1. Expression and purification of uAbs.** (a) Coomassie-stained SDS-PAGE analysis of E40-uAb, pE59-uAb, and EpE89-uAb and their corresponding DARPin counterparts E40, pE59, and EpE89 following expression in *E. coli* BL21(DE3) cells carrying a pET28-based plasmid corresponding to each of the constructs. Proteins were purified by Ni-NTA affinity chromatography with titers of ~30 mg protein per liter culture. Molecular weight ($M_W$) markers are indicated at left. Coomassie-stained gel is representative of at least three biological replicates. (b) Size exclusion chromatography (SEC) analysis of E40-uAb, pE59-uAb, and EpE89-uAb, and full-length human CHIP using a HiLoad Superdex 200 column. Peaks marked with asterisk (*) correspond to aggregated CHIP. CHIP eluted at a volume expected of a trimer or a dimer with a large water shell as assessed by comparison to elution of molecular weight standards.

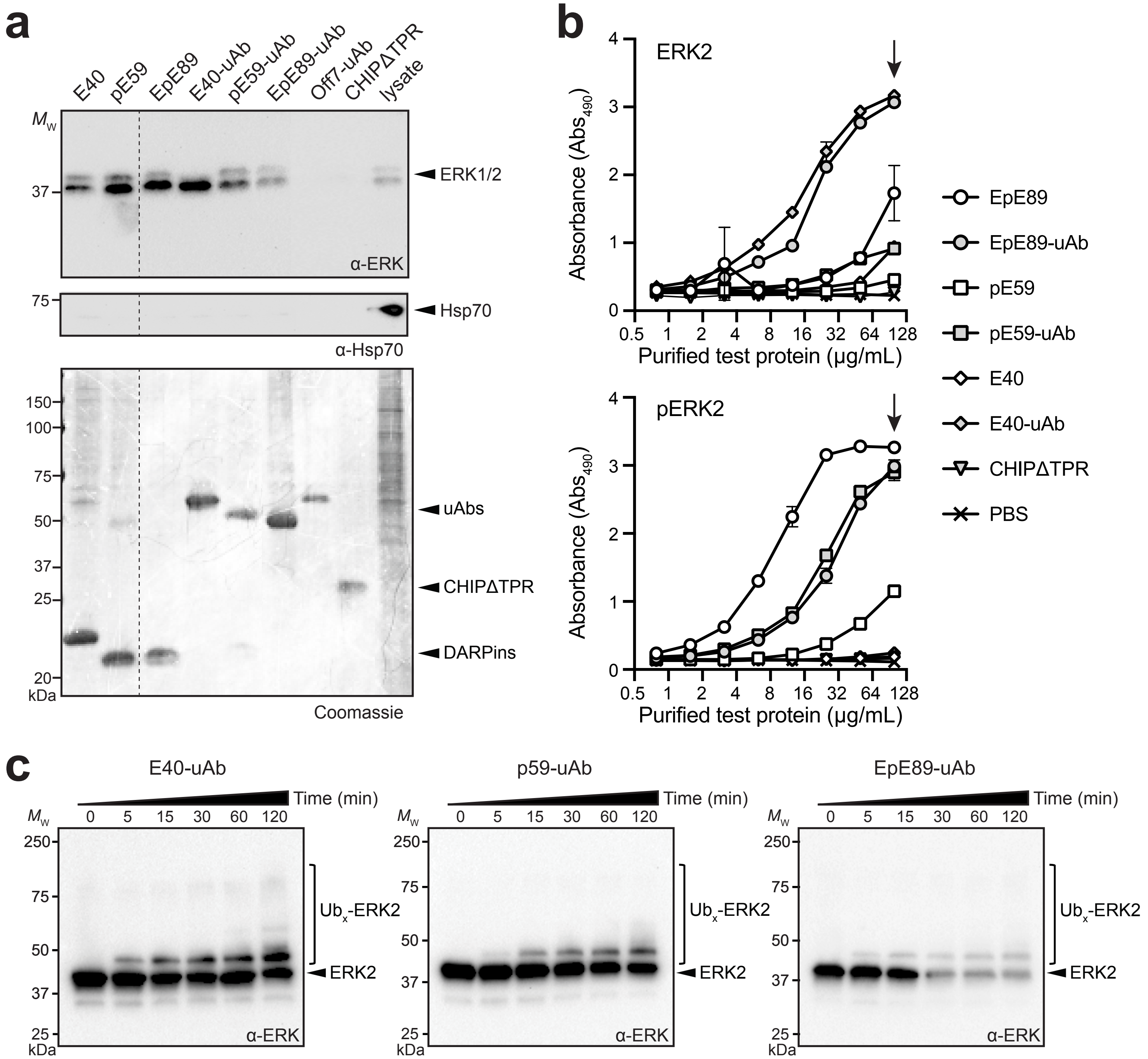


**Supplementary Figure 2. Binding and ubiquitination activity of ERK-directed uAbs.** (a) Affinity precipitation of ERK and pERK from HEK293T cells by E40-uAb, pE59-uAb, and EpE89-uAb and their corresponding DARPin counterparts E40, pE59, and EpE89. The off7-uAb and CHIPΔTPR constructs served as negative controls. Precipitated ERK2 and pERK were detected by immunoblot using a pan-ERK-specific antibody (α-ERK) that detects total ERK1/2. The ERK-directed uAbs and DARPins precipitated ERK1/2 forms from cell lysates, giving rise to the corresponding bands on a Western blot, whereas the off7-uAb and CHIPΔTPR controls did not. Immunoblotting with anti-Hsp70 (α-Hsp70) confirmed that Hsp70, a native substrate of human CHIP, was present in the lysate but not immunoprecipitated by any of the tested constructs including CHIP lacking its substrate binding domain (CHIPΔTPR). The presence of uAbs, DARPins, and control proteins was confirmed by Coomassie-stained SDS-PAGE analysis. (b) ELISA of purified uAbs, DARPins, and CHIPΔTPR against immobilized ERK2 (top) or pERK2 (bottom) as indicated. Buffer only (PBS) served as a negative control. Data are average of three biological replicates and error bars represent standard deviation of the mean. Arrow indicates the 100 µg/mL dilution that was reported in Figure 2c (c) *In vitro* ubiquitination of nonphosphorylated ERK2 in the presence of purified E40-uAb, pE59-uAb, or EpE89-uAb along with E1, E2, and ubiquitin (Ub). Samples were collected at indicated times and subjected to immunoblotting.

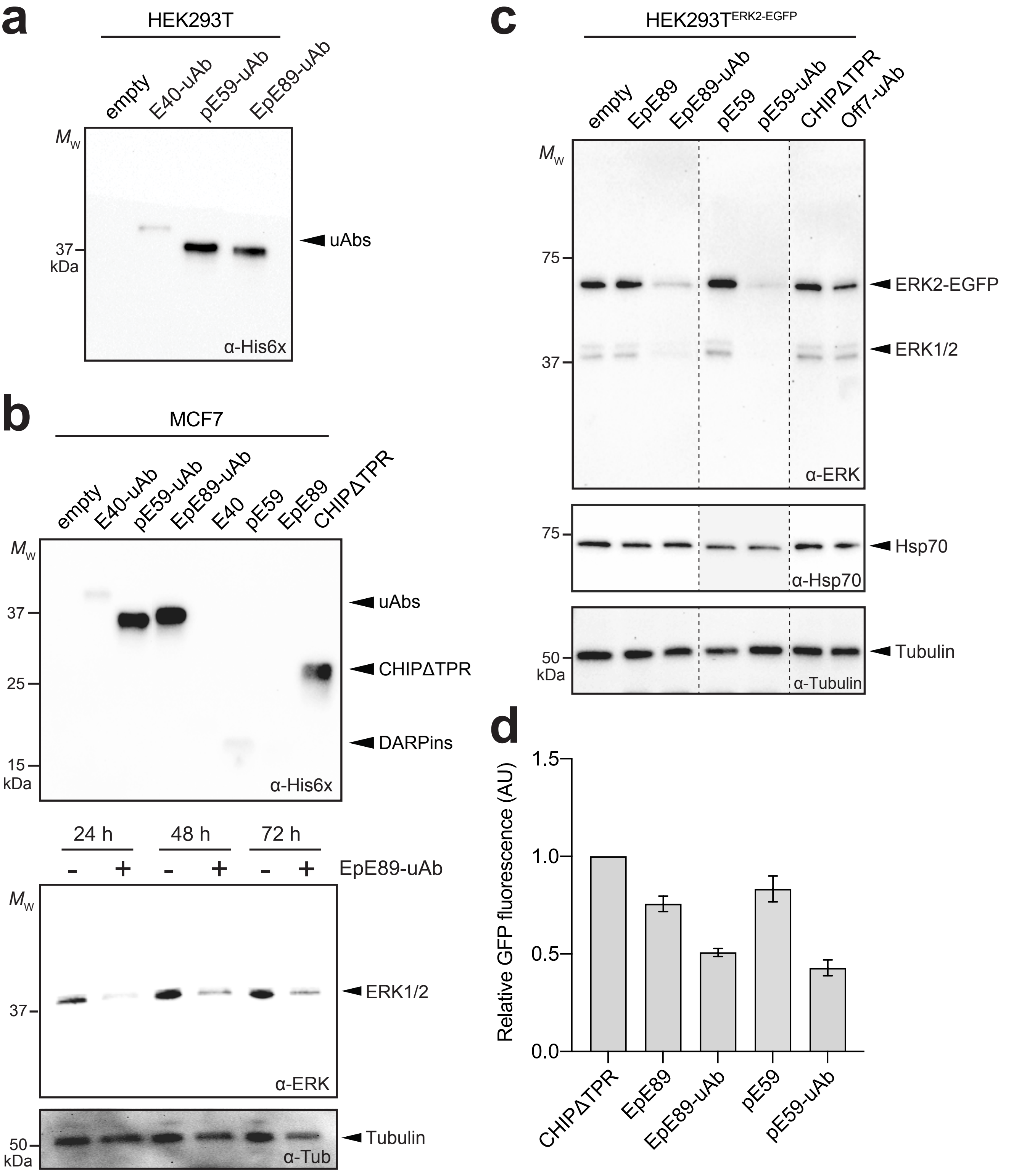


**Supplementary Figure 3. Expression and activity of ERK-targeting uAbs in living cells.** (a) Immunoblot analysis of extracts prepared from HEK293T cells transfected with empty pcDNA3 or pcDNA3 encoding each of the constructs indicated at 0.25 μg plasmid DNA per well. Cells were harvested 24 h post-transfection, after which extracts were prepared and subjected to immunoblotting. Blot was probed with anti-polyhistidine antibody (α-His6x) to detect uAb constructs. Lanes were normalized by total protein content. (b, top panel) Immunoblot analysis of extracts prepared from MCF7 cells transfected with empty pcDNA3 or pcDNA3 encoding each of the constructs indicated at 0.25 μg plasmid DNA per well. Blot was prepared exactly as in (a). (b, bottom panels) Immunoblot analysis of extracts prepared at the times indicated from MCF7 cells transfected with empty pcDNA3 (-) or pcDNA3 encoding EpE89-uAb (+). Lanes were normalized by total protein content. Blots were probed with pan-ERK antibody (α-ERK) to detect total ERK1/2 expression and equivalent loading was confirmed by probing with anti-β-tubulin (α-Tubulin). (c) Same as in (b) but with extracts prepared from HEK293T$^{ERK2-EGFP}$ cells transfected with empty pcDNA3 or pcDNA3 encoding each of the constructs indicated at 0.25 μg plasmid DNA per well. Lanes were normalized by total protein content and blots were probed with α-ERK, α-Hsp70 and α-Tubulin. For all blots, molecular weight ($M_W$) markers are indicated at left and results are representative of at least three biological replicates. (d) Flow cytometric analysis of EGFP fluorescence activity in select cells from (d) as indicated. Data are biological triplicates (three separately transfected wells) of the geometric mean fluorescence intensity (MFI) normalized to MFI measured for non-transfected HEK293T cells expressing ERK2-EGFP alone. Error bars represent standard deviation (SD) of the mean.

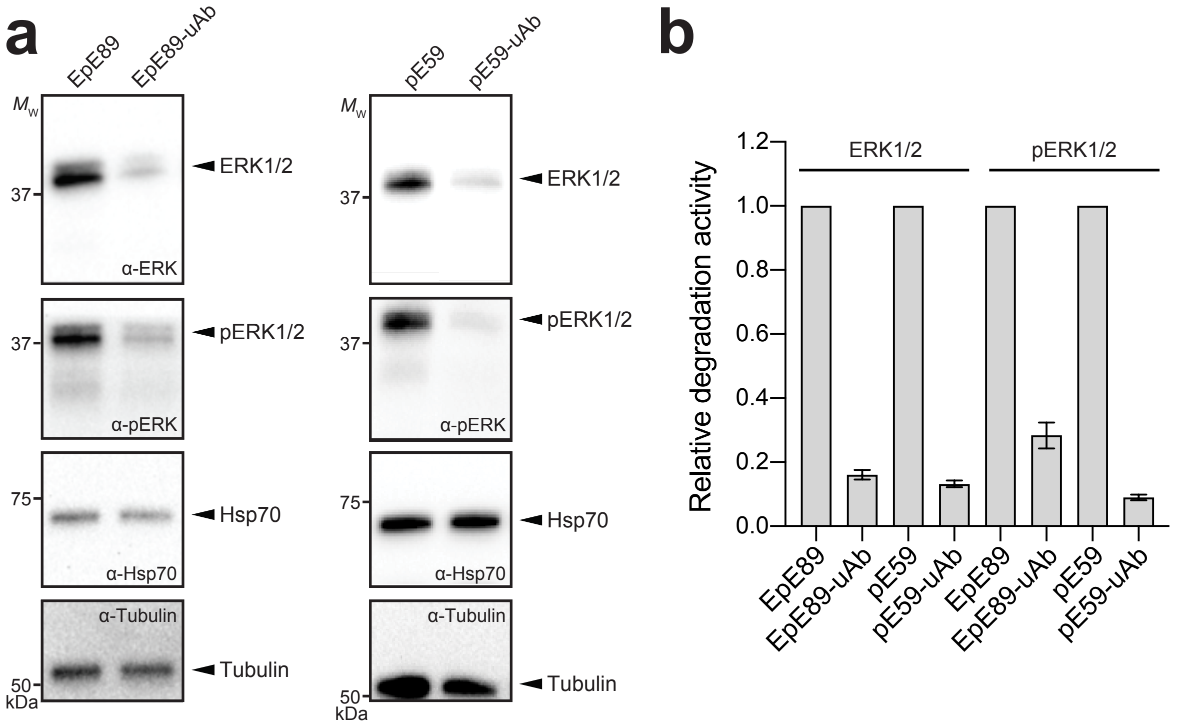


**Supplementary Figure 4. Silencing of pERK1/2 by EpE89 and pE59.** (a) Immunoblot analysis of extracts prepared from HEK293T cells transfected with pcDNA3 encoding EpE89, EpE89-uAb, pE59 and pE59-uAb at 0.25 µg plasmid DNA per well. Cells were harvested 24 h post-transfection, after which extracts were prepared and subjected to immunoblotting. Blots were probed with: pan-ERK antibody (α-ERK) to detect total ERK1/2 expression; anti-phospho-ERK antibody (α-pERK) to detect pERK1/2; and anti-Hsp70 antibody (α-Hsp70) to detect native CHIP substrate. Lanes were normalized by total protein content and equivalent loading was confirmed by probing with anti-β-tubulin (α-Tubulin). Molecular weight ($M_w$) markers are indicated at left. Results are representative of at least three biological replicates. (b) Relative quantitation of total ERK1/2 and pERK1/2 levels by densitometry analysis of α-ERK and α-pERK immunoblot images using ImageJ Software. Intensity data for uAb bands was normalized to band intensity for unfused DARPin control cases from three independent experiments. Error bars represent standard deviation of the mean.

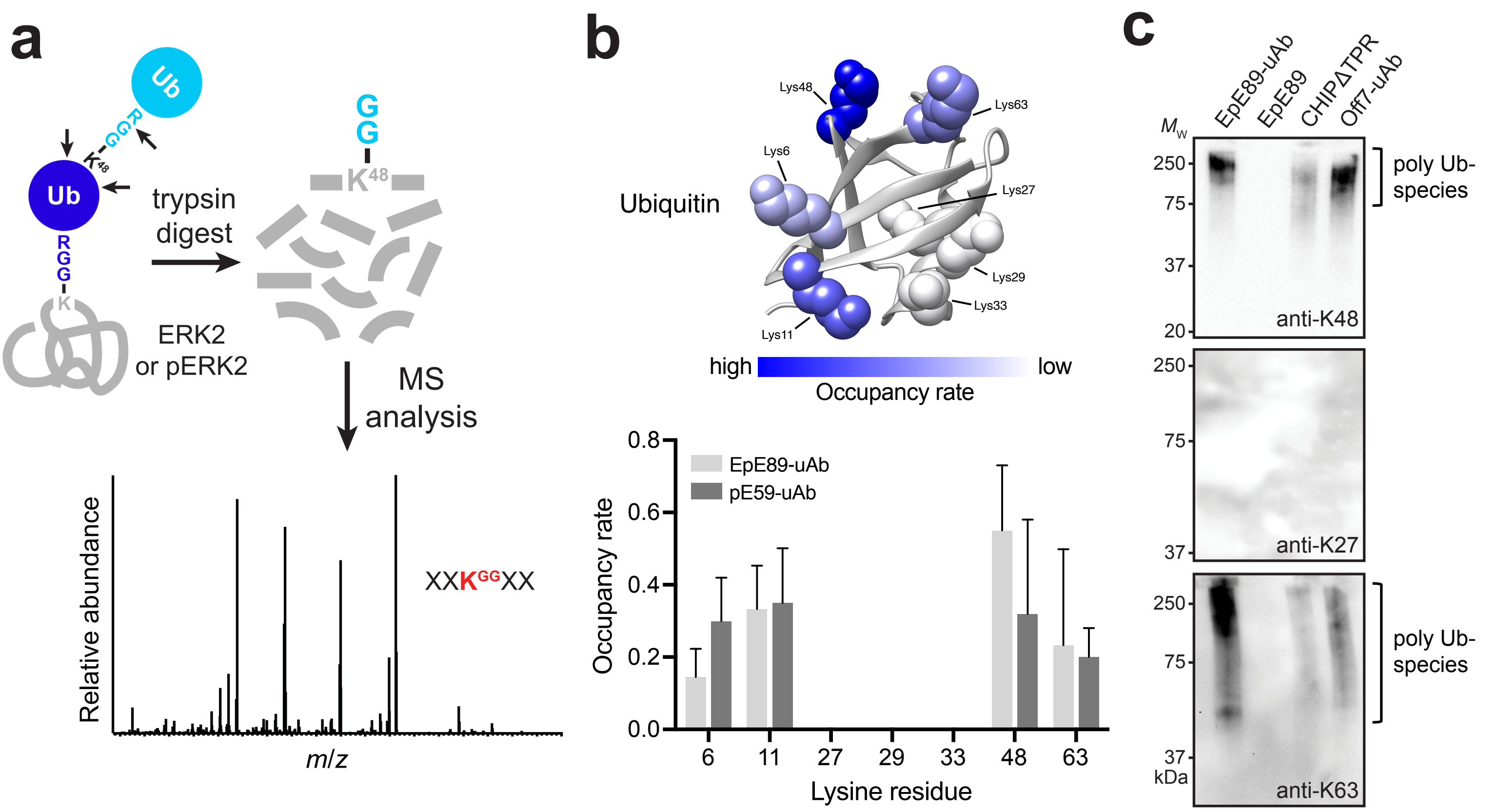


**Supplementary Figure 5. Determination of poly-Ub chain architecture by ubiquitin profiling.** (a) Schematic of the ubiquitin profiling method used to reveal poly-Ub chain architecture formed by uAb on ERK2/pERK2. Precise ubiquitination sites in Ub can be determined by mass spectrometry (MS) because of a characteristic mass shift caused by diglycine that is retained on lysine residues within Ub after trypsin digestion. Depending on the lysine residue in Ub that was modified with the Ub-Ub linkage, different linkage-specific signature peptides with characteristic masses are produced by trypsin digestion that can be readily detected by MS. (b) Occupancy rate of 'GG' modification of ubiquitin by EpE89-uAb and pE59-uAb determined by nanoLC-ESI-MS/MS analysis of in-gel trypsic digested samples. Data were generated by normalizing modified residue coverage with total residue coverage and averaging across six independent experiments, three with ERK2 and three with pERK2. Error bars represent the standard error of the mean (SEM). Crystal structure of human ubiqutin (white ribbon; PDB ID: 1UBQ rendered using PyMol Software) depicting seven lysine residues (spheres) colored according to the relative ubiquitination frequency of each lysine by EpE89-uAb, with dark blue indicating the most frequently modified sites and light blue indicating the least frequently modified sites. (c) Immunoblot analysis of extracts prepared from HEK293T cells transfected with pcDNA3 encoding each of the constructs indicated at 0.25 µg plasmid DNA per well. Cells were harvested 24 h post-transfection, after which extracts were prepared and subjected to immunoblotting. Blots were probed with: anti-K48, anti-K27, and anti-K63 antibodies to detect K48-, K27-, and K63-linked Ub products, respectively. Lanes were normalized by total protein content. Molecular weight ($M_W$) markers are indicated at left. Results are representative of at least three biological replicates.
